# Prevention of *Klebsiella pneumoniae* antibiotic resistance plasmid dissemination by prophylactic or simultaneous CRISPR-Cas9 treatment

**DOI:** 10.64898/2026.10.29.762760

**Authors:** Clàudia Morros-Bernaus, Nia Feakes, Urszula Oko, Keira Cozens, Stineke van Houte, David Sünderhauf

## Abstract

Antimicrobial resistance (AMR) is one of the most prevailing challenges of the 21^st^ century, driven in part by pathogens such as *Klebsiella pneumoniae* which frequently disseminate plasmids carrying AMR genes within and across species. Mobile CRISPR-Cas9 tools can limit the spread of AMR plasmids through selective targeting but suffer efficacy issues when faced with ecological realism.

Here, we assembled a simple three-strain community to assay the effectivity of mobile CRISPR-Cas9 in preventing dissemination of a clinical *K. pneumoniae bla*_CTX-M-15_ plasmid from its native host. We applied an *Escherichia coli* donor carrying a mobile CRISPR-Cas9 plasmid and entirely prevented transfer of the *K. pneumoniae* AMR plasmid into a focal, plasmid-free, *E. coli* recipient. Under experimental conditions favouring CRISPR-Cas9 plasmid transfer, AMR plasmid uptake was blocked regardless of whether the CRISPR-Cas9 plasmid spread prophylactically or simultaneously with the *K. pneumoniae* AMR plasmid.

Our work shows that mobile CRISPR-Cas9 tools can overcome barriers to successfully prevent dissemination of AMR plasmids. Prophylactic application is not essential, and AMR plasmid transfer can even be suppressed when AMR plasmids spread simultaneously with mobile CRISPR-Cas9 tools.

## 1. INTRODUCTION

*Klebsiella pneumoniae* is a member of the *Enterobacteriaceae* and commensally associated with environmental and animal microbiomes (Lee et al., 2021; Ribeiro et al., 2022). In humans, *K. pneumoniae* most frequently colonizes gut microbiomes and can opportunistically become pathogenic and cause disease such as pneumonia, urinary tract, and bloodstream infections (Martin & Bachman, 2018; Wyres et al., 2020).

*K. pneumoniae* presents a global threat to human health and is on the World Health Organization list of critical priority bacterial pathogens (World Health Organization, 2024), in particular due to its propensity to acquire genes conferring resistance to broad-spectrum beta-lactam antibiotics including carbapenems and third-generation cephalosporins (De Oliveira et al., 2020). Many of these antimicrobial resistance (AMR) genes are located on mobile genetic elements, most notably plasmids. *K. pneumoniae* plasmid-mediated AMR spread is highly important in the mobilisation of AMR genes from environmental into clinical bacteria (Wyres & Holt, 2018); usually into other strains of *K. pneumoniae* (Pedersen et al., 2020). Plasmid mobilisation into other species is less common, but does occur (León-Sampedro et al., 2021; Lindblom et al., 2019; Sandegren et al., 2012).

Dissemination of AMR plasmids within patients is particularly problematic for two reasons: Firstly, *K. pneumoniae* may disseminate AMR plasmids into pathogenic *Enterobacteriaceae* such as *Escherichia coli* (Bidet et al., 2005), compromising otherwise successful antibiotic treatment. Secondly, plasmid dissemination into other gut commensals can give rise to novel plasmid hosts (Jeong et al., 2015; Schjørring et al., 2008; Xavier et al., 2019). Highly successful plasmid-host combinations in particular have the potential to become important new plasmid sources to the wider microbial community, causing community-level plasmid persistence where plasmids may otherwise become lost (Alonso-del Valle et al., 2021; Li et al., 2020). Therefore, preventing onwards dissemination of AMR plasmids from *Klebsiella* may be a means of reducing the overall AMR burden within patients or other environments.

CRISPR-Cas technologies offer a promising strategy for reducing plasmid dissemination. Due to its high programmability and sequence-specificity defined by its guide RNA, CRISPR-Cas9 can selectively target plasmids carrying a trait of interest, such as a clinically relevant AMR gene (Bikard et al., 2014; Citorik et al., 2014). We previously generated the broad host-range conjugative plasmid pKJK5::csg, engineered to encode the Cas9 nuclease, a single guide RNA, and GFP. Use of this CRISPR-Cas9 plasmid demonstrated selective removal of an AMR cloning vector from *Escherichia coli* and prevention of uptake of AMR vectors in a range of strains belonging to the orders *Pseudomonadales* and *Enterobacterales* (Sünderhauf et al., 2023).

When targeting larger, conjugative AMR plasmids which encode competitive traits such as toxin-antitoxin (TA) systems, we found that CRISPR-Cas9 is more effective when protecting a host from AMR plasmid uptake. In contrast, CRISPR-Cas9 can be selected against when trying to invade a new host where it displaces an established AMR plasmid (Sünderhauf et al., 2026). Ecological realism may also lead to lower conjugation rates, which similarly reduced effectivity of CRISPR-Cas-directed plasmid curing, whilst CRISPR-Cas remained a strong barrier against AMR plasmid uptake (Upreti et al., 2024). In line with this, we hypothesise that the use of mobile CRISPR-Cas9 tools to prevent dissemination of AMR plasmids into antibiotic sensitive hosts could be more effective than displacing plasmids from their resident host, but this idea remains untested. Specifically, we hypothesise that preventing dissemination of *K. pneumoniae* plasmids to other bacterial species is an especially effective application of CRISPR-Cas9-based approaches.

Here, we carry out mating experiments to disrupt dissemination of a clinical *K. pneumoniae bla*_CTX-M-15_ plasmid to *E. coli*. We achieved this by allowing a CRISPR-Cas9 plasmid, hosted in *E. coli* K12, to spread into *E. coli* DH5α before exposure to *K. pneumoniae*. Further, under mating conditions that favour transfer of the CRISPR-Cas9 plasmid, AMR plasmid dissemination was also prevented when *E. coli* DH5α was exposed to *K. pneumoniae* and the CRISPR-Cas9 plasmid simultaneously. High transfer of the CRISPR-Cas9 plasmid reduced overall *K. pneumoniae* burden, highlighting multiple benefits of applying mobile CRISPR-Cas9 plasmids to prevent *K. pneumoniae* AMR plasmid dissemination in microbial communities.

## 2. RESULTS

### 2.1 Genomic characterisation of clinical isolate MA48971.2

Clinical isolate *K. pneumoniae* MA48971.2, obtained from a patient at University Medical Center, Utrecht (Netherlands) in 2018, was sequenced and characterised by hybrid assembly. We found that the isolate carried four plasmids and belonged to sequence type (ST) 353 (***Table 1***), which has been linked to disease (Böhne et al., 2026; Gelalcha et al., 2024). The strain was found to carry eight resistance genes, including ESBLs and *bla*_CTX-M-15_, which confers resistance to cefotaxime. The lack of carbapenemases or colistin resistance gave MA48971.2 a moderate Kleborate resistance score of 1. Importantly, four of its resistance genes were found on a single ∼200 kb IncFII/IncFIB plasmid (pDSU25-1.1F, ***Table 1***), highlighting strong potential for AMR gene dissemination. In terms of virulence, the isolate was found to chromosomally carry the yersiniabactin locus and thereby had a Kleborate virulence score of 1. As a *bla*_CTX-M-15_-positive *K. pneumoniae* ST353 strain, the AMR profile of our focal clinical isolate is representative of other *K. pneumoniae* of the same clade reported throughout literature (Jiang et al., 2026).

**Table 1:** Genomic characterisation of *K. pneumoniae* isolate MA48971.2. non-mob – non-mobile as identified by MOB-typer, see *Methods*.

|  | Chromosome | pDSU25-1.1F | pDSU25-1.2 | pDSU25-1.3col | pDSU25-1.4 |
| --- | --- | --- | --- | --- | --- |
| **Size** | 5272866 bp | 201056 bp | 4406 bp | 4258 bp | 2821 bp |
| **Replicon Types** | - | IncFIB, IncFII, rep_cluster_2183 | - | ColRNAI_rep_cluster_1987 | - |
| **Predicted Mobility** | - | conjugative | non-mob; putative oriT at 93% identity | non-mob; putative oriT at 79% identity (across 19% sequence) | non-mob; putative oriT at 93% identity (across 68% sequence) |
| **AMR Genes** | OqxB, OqxA, blaSHV-187, fosA | bla<sub>TEM-1B</sub>, bla<sub>CTX-M-15</sub>, aac(6')-Ib-cr, bla<sub>OXA-1</sub> | - | - | - |
| **TA systems** | - | pemIK, higAB, tacAT (2x), vagCD | - | - | - |
| **Defense Systems** | PD-T4-3, RM_Type_I (5x), Taranis, Rst_TIR-NLR, Other_Type_IV, Rst_RT-nitrilase-Tm, AbiE (2x) | MazEF (2x), TgvAB (2x), RM_Type_I (3x), Mok_Hok_Sok | - | PD-T4-4 (2x) | - |

### 2.2 Condition-dependent plasmid mobilisation readily occurs into the focal *E. coli* DH5α recipient

Before investigating the protective effect of CRISPR-Cas9 against the spread of *K. pneumoniae* AMR plasmids, we assessed individual plasmid transfer dynamics to identify optimal experimental conditions for mobile CRISPR-Cas9 application. We first measured transfer of AMR plasmid pDSU25-1.1F into focal recipient *E. coli* DH5α, making use of this straińs chromosomal streptomycin resistance tag (see Methods). Since plasmid mobility is impacted by media type (Sheppard et al., 2020) we performed mating experiments in both solid-agar and liquid-broth conditions. Selective plating after 48h mating revealed that transfer of pDSU25-1.1F from *K. pneumoniae* to *E. coli* DH5α was significantly higher in liquid-broth mating compared to solid-agar (gamma GLM, emmeans contrast, *p* <0.001) (***Figure 1A***), achieving a conjugation efficiency of 2.2 ± 0.52 x 10^-2^ transconjugants per recipients.

**Figure 1:**
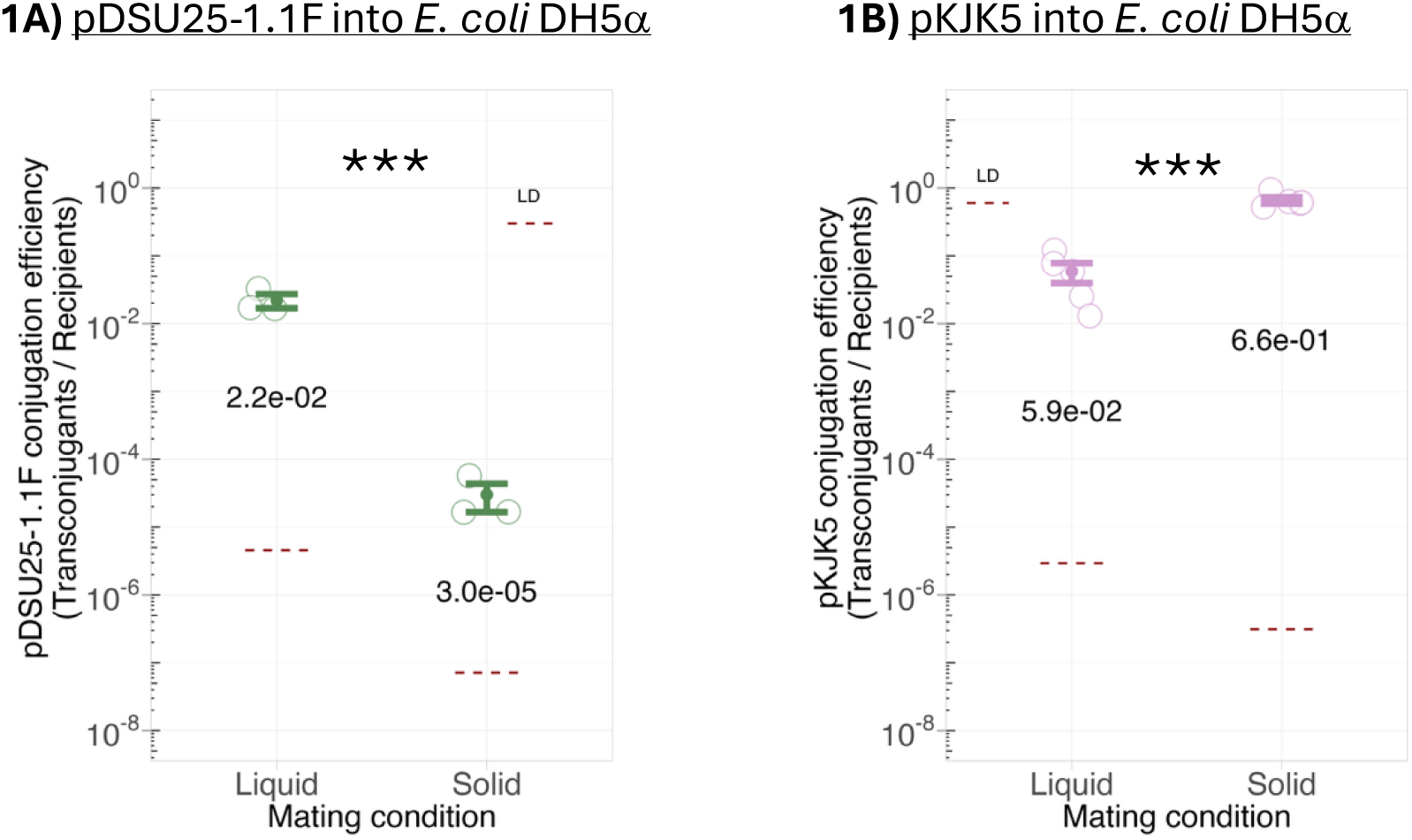
Transfer of pDSU25-1.1F and of pKJK5::csgc[NT] into *E. coli* DH5α in one-to-one matings. **A)** Transfer of pDSU25-1.1F from *K. pneumoniae* MA48971.2 to *E. coli* DH5α in liquid-broth and solid-agar mating. Plasmid transfer is shown as conjugation efficiency, which is calculated as transconjugants (*E. coli* DH5α + pDSU25-1.1F) divided by recipients (overall *E. coli* DH5α population). *** *p* = 0.0003 as assessed by gamma GLM, emmeans contrast. Mean and standard error are shown as point and bars, individual replicates are plotted as circles. The limit of detection (LD) of the assay is plotted as a dashed red line. LD was determined based on plating volume and dilution factors and was set to 0.5 transconjugants detected in the non-diluted sample. Since LD is different for each individual replicate, the highest LD within each treatment is plotted. **B)** Transfer of pKJK5::csgc[NT] from *E. coli* K12 to *E. coli* DH5α in liquid-broth and solid-agar mating. *** *p* = 0.0003 as assessed by gamma GLM, emmeans contrast.

Alongside this, we measured transfer of CRISPR-Cas9 plasmid pKJK5::csgc (*cas9, sgRNA, gfp, cat*) from its donor *E. coli* K12 strain (which represses pKJK5’s *gfp* expression; see Methods) into the same plasmid-free *E. coli* DH5α focal recipient. For this experiment, we used pKJK5::csgc[NT], which is used as a non-targeting control and carries a guide RNA directed against a 20-nucleotide sequence absent from the genomes of all three bacterial strains. In contrast to *K. pneumoniae* plasmid transfer, transfer of pKJK5::csgc[NT] from *E. coli* K12 to *E. coli* DH5α was significantly higher in solid-agar than in liquid-broth mating (gamma GLM, emmeans contrast, *p* <0.001) (***Figure 1B***), achieving values of up to 6.6 ± 0.73 x 10^-1^ transconjugants per recipients.

Overall, this demonstrates that transfer of both pDSU25-1.1F and of pKJK5::csgc[NT] into the recipient *E. coli* DH5α strain occurs and is strongly influenced by mating conditions. After liquid-broth mating, pKJK5::csgc[NT] and pDSU25-1.1F were transferred at a similar frequency of ∼10^-2^ transconjugants per recipients. Therefore, under these conditions, we expect *E. coli* DH5α to be equally likely to acquire either of the plasmids if both are present in a community at similar frequencies. In contrast, under solid-agar mating conditions, we expect *E. coli* DH5α to be far more likely to acquire pKJK5::csgc[NT] (conjugation efficiency ∼10^-1^ transconjugants per recipients) than pDSU25-1.1F (conjugation efficiency ∼10^-5^ transconjugants per recipients). Consequently, in solid-agar mating the *bla*_CTX-M-15_-targeting version of the CRISPR-Cas9 plasmid (pKJK5::csgc[*bla*_CTX-M-15_]) may effectively function as a plasmid ‘vaccine’ by establishing in *E. coli* DH5α and protecting against pDSU25-1.1F acquisition.

### 2.3 Prophylactic application of *E. coli* K12 + pKJK5::csgc[*bla*_CTX-M-15_] protects *E. coli* DH5α from pDSU25-1.1F uptake

To assess whether the CRISPR-Cas9 plasmid can act as a plasmid ‘vaccine’ to block pDSU25-1.1F uptake, we combined the three strains to form a simple synthetic community: (i) focal recipient *E. coli* DH5α, susceptible to plasmid acquisition, (ii) CRISPR-Cas9 plasmid donor *E. coli* K12 and (iii) clinical isolate *K. pneumoniae* MA48971.2, carrying its *bla*_CTX-M-15_ plasmid pDSU25-1.1F **(*Figure 2A*).** To measure the impact of CRISPR-Cas9 targeting, *E. coli* K12 carried either pKJK5::csgc[*bla*_CTX-M-15_], which contains a guide RNA targeting the AMR gene *bla*_CTX-M-15_, or pKJK5::csgc[NT] with its non-targeting guide. This experimental setup allowed us to assay the specific effect of CRISPR-Cas9 targeting. To measure the impact of pKJK5::csgc presence in the community (independent of CRISPR-Cas9 effects), we included an additional “no plasmid” treatment where *E. coli* K12 remained empty.

**Figure 2:**
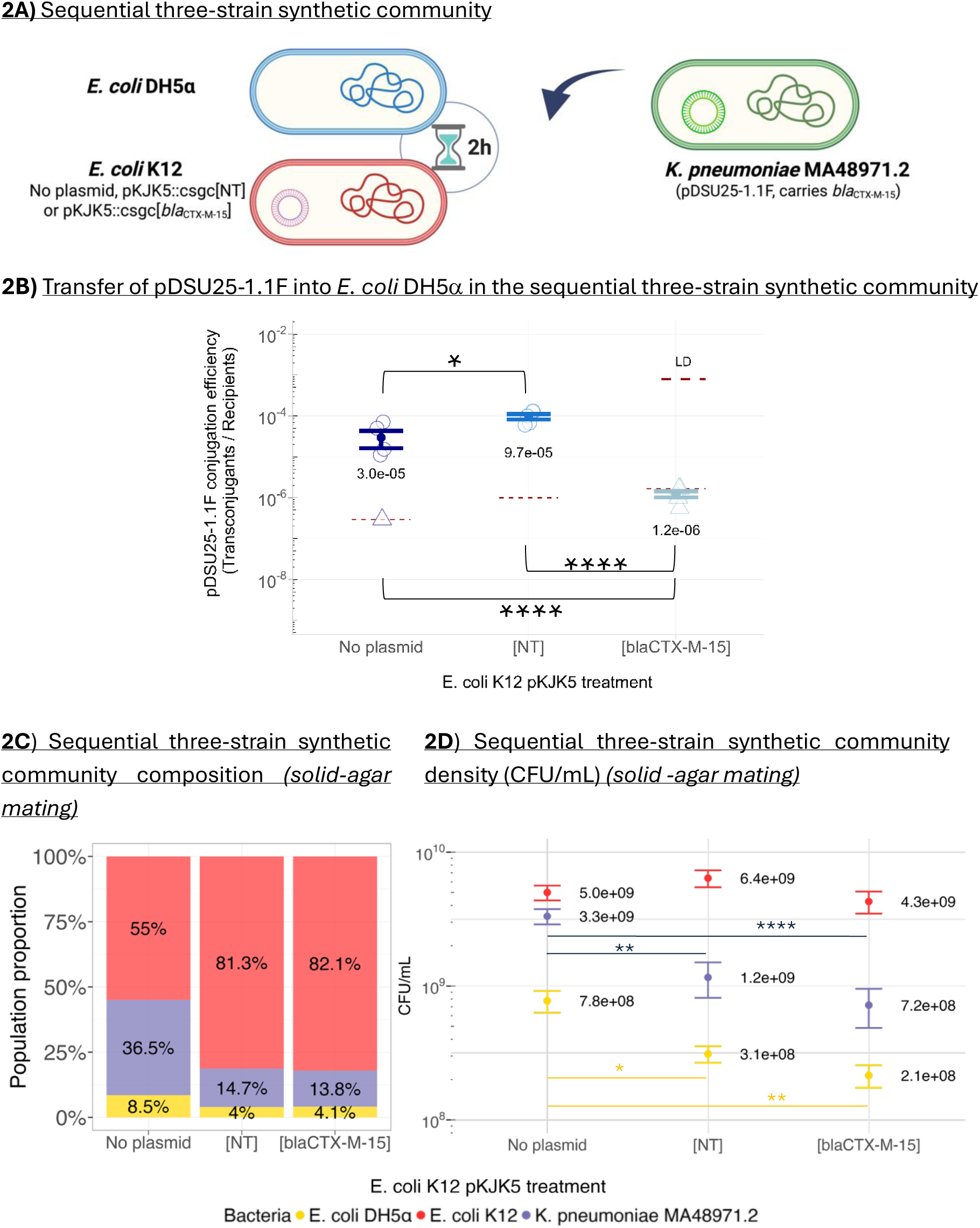
Representation of the sequential three-strain synthetic community (**A**), CRISPR-Cas9 protection against uptake of AMR plasmid and population dynamics in solid-agar mating (**B-D**). **2A)** Representation of the sequential three-strain synthetic community formed by *K. pneumoniae* MA48971.2, *E. coli* K12 and *E. coli* DH5α. *K. pneumoniae* carries ∼200 kb IncFII/IncFIB plasmid (pDSU25-1.1F) which carries *bla*_CTX-M-15_ and is added 2h after mating between the two *E. coli* strains. *E. coli* K12 carries either no plasmid, pKJK5::csgc[NT] or pKJK5::csgc[*bla*_CTX-M-15_]. **2B)** Transfer of the *K. pneumoniae* pDSU25-1.1F into *E. coli* DH5α in the sequential three-strain community as assessed by selective plating, shown as conjugation efficiency, which is calculated as transconjugants (*E. coli* DH5α + pDSU25-1.1F) divided by recipients (overall *E. coli* DH5α population). Mean and standard error are shown as point and bars and individual replicates are plotted as circles. In those cases where pDSU25-1.1F-carrying transconjugants were not observed, the limit of detection (LD) of the assay is plotted instead (denoted by triangles). LD was determined based on plating volume and dilution factors and was set to 0.5 transconjugants detected in the non-diluted sample. Since LD is different for each individual replicate, the highest LD within treatment is plotted as a dashed red line. \**p*<0.05, \*\*\*\**p*<0.0001 as assessed by gamma GLM, emmeans contrast Tukey-adjusted. **2C-D)** Community composition (**C**) and bacterial densities (**D**) across *E. coli* K12 treatments. *E. coli* K12 is shown in red, *E. coli* DH5α in yellow and *K. pneumoniae* in purple. **C)** Bacterial community composition. Means from 5 replicates are plotted across treatments. Plasmid treatment significantly affected bacterial community composition (MANOVA, V = 0.82, F (4,24) = 4.18, p = 0.01). **D)** Bacterial density of the respective strains (CFU/mL). Mean and standard error are shown as point and bars. \**p*<0.05, \*\**p*<0.01, \*\*\*\**p*<0.0001 as assessed by gamma GLM, emmeans contrast Tukey-adjusted. Other within-strain comparisons remained non-significant.

We combined the three strains sequentially, allowing *E. coli* K12 and *E. coli* DH5α to mate in solid-agar conditions for two hours, after which existing mating pairs were disrupted and *K. pneumoniae* was added **(*Figure 2A*)**. This initial two-hour mating between *E. coli* K12 and *E. coli* DH5α allowed sufficient time for transfer of pKJK5::csgc[*bla*_CTX-M-15_] into *E. coli* DH5α (conjugation efficiency up to 8.3 ± 2.04 x10^-1^ transconjugants per recipients; ***Supplementary figure 1***).

After addition of *K. pneumoniae* and solid-agar mating of the microbial community for an additional 48 hours, we observed AMR plasmid uptake block in *E. coli* DH5α attributable to pKJK5::csgc[*bla*_CTX-M-15_]. The *E. coli* DH5α uptake of pDSU25-1.1F remained entirely underneath the limit of detection and was therefore significantly reduced by at least ∼2 orders of magnitude in the presence of *E. coli* K12 + pKJK5::csgc[*bla*_CTX-M-15_] compared to the non-targeting pKJK5::csgc[NT] (gamma GLM, emmeans contrast Tukey-adjusted, *p* <0.0001), or the no plasmid *E. coli* K12 treatment (*p* <0.0001) (***Figure 2B***). Most *E. coli* DH5α + pDSU25-1.1F transconjugants recovered in the presence of *E. coli* K12 + pKJK5::csgc[NT] were GFP+, revealing the acquisition of both pDSU25-1.1F and pKJK5::csgc[NT]. However, such co-existence was disrupted in presence of pKJK5::csgc[*bla*_CTX-M-15_], consistent with CRISPR-Cas9-directed cleavage of pDSU25-1.1F.

As expected, pKJK5::csgc[*bla*_CTX-M-15_] was a very strong barrier to pDSU25-1.1F uptake in *E. coli* K12 too and entirely prevented invasion by the AMR plasmid (**S*upplementary figure* 2**).

Overall, these results are in line with the higher transfer of pKJK5::csgc than of pDSU25-1.1F into *E. coli* DH5α observed in the previous one-to-one matings (***Figure 1*)**. As expected, a parallel experiment under liquid-broth mating conditions showed reduced transfer of pKJK5::csgc into *E. coli* DH5α (∼10^-4^ transconjugants per recipients) (***Supplementary figure 1***), leading to no observable reduction in pDSU25-1.1F dissemination in presence of pKJK5::csgc[*bla*_CTX-M-15_] (***Supplementary figure 3A***).

Altogether, this data demonstrates the probiotic profile of *E. coli* K12 + pKJK5::csgc[*bla*_CTX-M-15_] when administered prophylactically in a microbial community, dependent on efficient pKJK5::csgc mobilisation. The presence of *E. coli* K12 + pKJK5::csgc[*bla*_CTX-M-15_] protected both itself and *E. coli* DH5α from pDSU25-1.1F uptake.

### 2.4 pKJK5 carriage reduces *K. pneumoniae* burden in the three-strain synthetic community

After confirming a CRISPR-Cas9-mediated protective effect in the recipient *E. coli* DH5α strain, we also analysed whether the respective *E. coli* K12 treatments had an impact on the overall synthetic community composition. We observed a significant shift in the bacterial community composition between *E. coli* K12 treatments (MANOVA, V = 0.82, *F* (4,24) = 4.18, *p* = 0.01) (***Figure 2C***). This was driven by a significant 63% and 78% reduction of the *K. pneumoniae* burden in both the pKJK5::csgc[NT] (gamma GLM, emmeans contrast Tukey-adjusted *p* = 0.003) and the pKJK5::csgc[*bla*_CTX-M-15_] (*p* <0.0001) treatments respectively, compared to the no plasmid treatment (***Figure 2D***). We could not identify a direct impact of CRISPR-Cas9 targeting on *K. pneumoniae* abundance; an observed 40% reduction between [NT] and [*bla*_CTX-M-15_] treatments remained non-significant (*p* = 0.3) (***Figure 2D***).

In contrast, under liquid-broth mating conditions (where pDSU25-1.1F dissemination could not be prevented; ***Supplementary figure 3A***), community composition also remained unchanged (*p* = 0.6) (***Supplementary figure 3B-C***).

Overall, these results indicate that the presence of pKJK5::csgc itself, independent of CRISPR-Cas9 targeting activity, can reduce *K. pneumoniae* strain abundance under mating conditions that allow high transfer of pKJK5::csgc within the synthetic community.

### 2.5 Application of pKJK5::csgc[*bla*_CTX-M-15_] simultaneous to *K. pneumoniae* exposure conserves protective effects

After observing how prophylactic application of *E. coli* K12 + pKJK5::csgc[*bla*_CTX-M-15_] protects the synthetic community against pDSU25-1.1F uptake and reduces the burden of the AMR-carrying *K. pneumoniae* strain, we considered that prophylactic administration of the CRISPR-Cas9 donor may not always be feasible in realistic applications. We therefore next assessed whether such results could also be achieved upon simultaneous exposure of *E. coli* DH5α to both *K. pneumoniae* MA48971.2 and to *E. coli* K12.

This experiment recapitulated the previously observed effects: under solid-agar mating conditions, a significant reduction of pDSU25-1.1F uptake in *E. coli* DH5α was observed in the presence of *E. coli* K12 + pKJK5::csgc[*bla*_CTX-M-15_], compared to [NT] (gamma GLM, emmeans contrast Tukey-adjusted *p* <0.0001) or the no plasmid treatment (*p* <0.0001) (***Figure 3A***). Again, no *E. coli* DH5α pDSU25-1.1F transconjugants were detected in the presence of *E. coli* K12 carrying pKJK5::csgc[*bla*_CTX-M-15_], while all the *E. coli* DH5α + pDSU25-1.1F transconjugants recovered in the presence of *E. coli* K12 + pKJK5::csgc[NT] were GFP+, revealing the co-existence of both pDSU25-1.1F and pKJK5::csgc[NT] in the cells. These results demonstrate that the pKJK5::csgc vaccination effect can be achieved through simultaneous introduction of all members of the synthetic community when mating conditions allow for high transfer of pKJK5::csgc[*bla*_CTX-M-15_].

**Figure 3:**
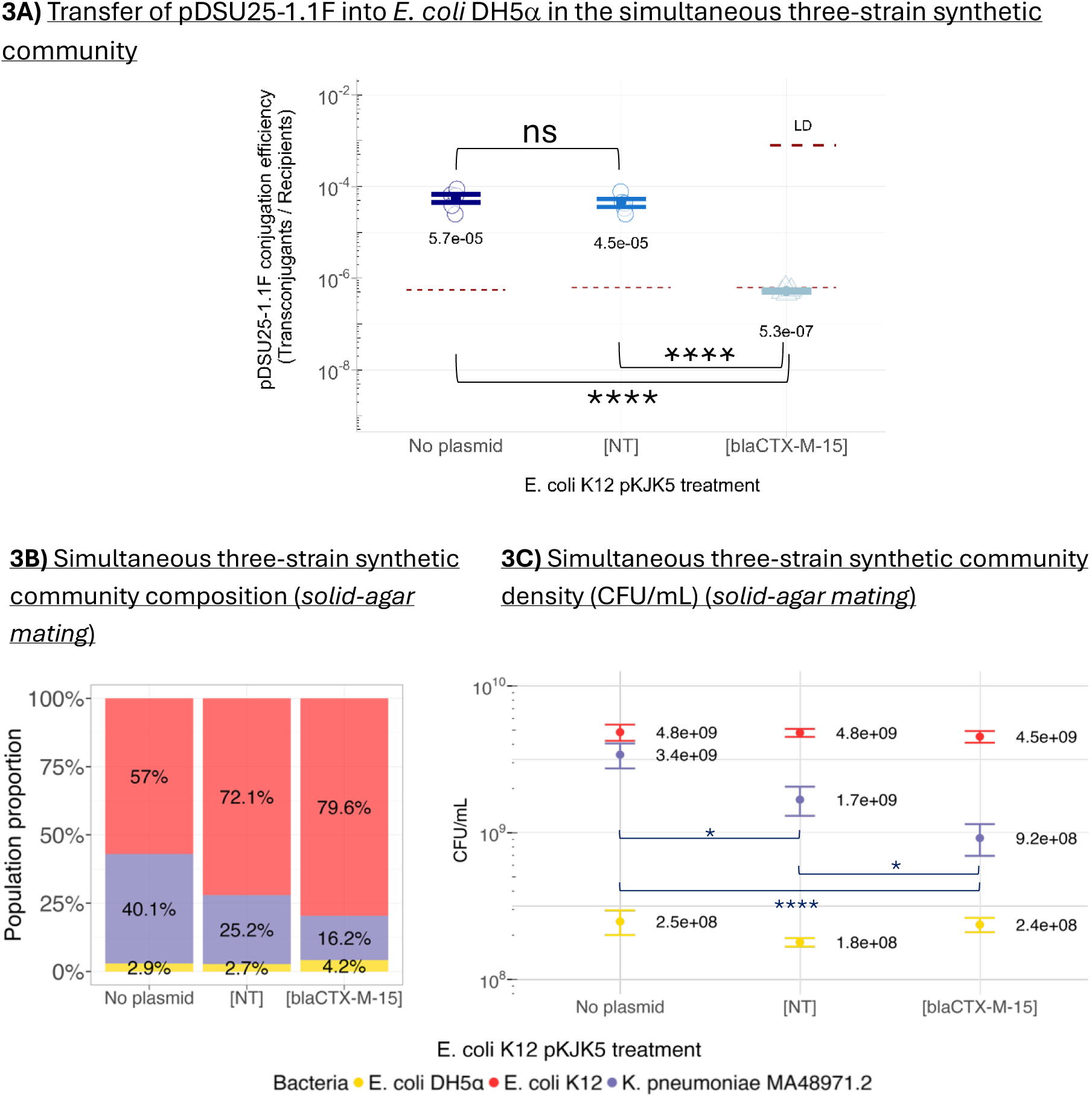
CRISPR-Cas9 protection against uptake of AMR plasmid and population dynamics in the solid-agar mating of the simultaneously assembled three-strain synthetic community (**A-C**)**. 3A)** Transfer of the *K. pneumoniae* plasmid pDSU25-1.1F into *E. coli* DH5α in the three-strain synthetic community, (as in Figure 2A, but with simultaneous strain introduction) is shown as conjugation efficiency, which is calculated as transconjugants (*E. coli* DH5α + pDSU25-1.1F) divided by recipients (overall *E. coli* DH5α population). Mean and standard error are shown as point and bars and individual replicates are plotted as circles. In those cases where pDSU25-1.1F-carrying transconjugants were not observed, the limit of detection (LD) of the assay is plotted instead (denoted by triangles). LD was determined based on plating volume and dilution factors and was set to 0.5 transconjugants detected in the non-diluted sample. Since LD is different for each individual replicate, the highest LD within treatment is plotted as a dashed red line. N.s. not-significant; \*\*\*\**p*<0.0001 as assessed by gamma GLM, emmeans contrast Tukey-adjusted. **B-C)** Synthetic community composition (**B**) and bacterial densities (**C**) across *E. coli* K12 treatments. *E. coli* K12 is shown in red, *E. coli* DH5α in yellow and *K. pneumoniae* in purple. **3B)** Bacterial community composition. Means from 5 replicates are plotted across treatments. Plasmid treatment significantly affected bacterial community composition (MANOVA, V= 0.67, *F* (4,24) = 3.07, *p* = 0.036). **3C)** Bacterial density of the respective strains (CFU/mL), mean and standard error are shown as point and bars. \**p*<0.05, \*\*\*\**p*<0.0001 as assessed by gamma GLM, emmeans contrast Tukey-adjusted. Other within-strain comparisons remained non-significant.

Additionally, the synthetic community composition was also significantly impacted by the different *E. coli* K12 treatments (*p* = 0.036) (***Figure 3B***). This effect was entirely driven by a significant 50% and 72% reduction of the *K. pneumoniae* burden between the no plasmid and the pKJK5::csgc[NT] (*p* = 0.04) or pKJK5::csgc[*bla*_CTX-M-15_]) (*p*<0.0001) treatments, respectively (***Figure 3C***). Unlike after sequential mating, after simultaneous community member introduction CRISPR-Cas9 targeting also decreased *K. pneumoniae* proportion: a statistically significant 45% reduction of the *K. pneumoniae* population was detected between treatments (from 1.7 ± 0.38 x 10^9^ in pKJK5::csgc[NT] to 9.2 ± 3.8 x 10^8^ in pKJK5::csgc[*bla*_CTX-M-15_]; *p* = 0.03; ***Figure 3C***).

As in the sequential experiment, solid-agar mating conditions were necessary to observe these effects and a parallel liquid-broth mating experiment, which saw low levels of pKJK5::csgc transfer, led to no protection from AMR plasmid uptake or reduction of *K. pneumoniae* burden (***Supplementary figure 4***).

Overall, in this study we developed a three-strain synthetic community that contains both a clinical AMR plasmid (*K. pneumoniae,* pDSU25-1.1F) and a CRISPR-Cas9 plasmid (*E. coli* K12, pKJK5::csgc) and we assessed the transfer of the respective plasmids and the impact of CRISPR-Cas9 targeting. We showed that under conditions that allow high transfer of pKJK5::csgc, as achieved in solid-agar mating, the presence of *E. coli* K12 + pKJK5::csgc[*bla*_CTX-M-15_] protects both itself and a focal recipient *E. coli* DH5α from pDSU25-1.1F uptake. Additionally, the presence of pKJK5::csgc itself, even in absence of CRISPR-Cas9 targeting, also reduces the burden of the AMR-containing strain and such reduction can be maximised by CRISPR-Cas9 targeting. Finally, we showed that the protective effect of *E. coli* K12 + pKJK5::csgc[*bla*_CTX-M-15_] within the synthetic community is observed regardless of whether it is introduced prophylactically before the exposure to the AMR-carrying strain or simultaneously with it.

## 3. DISCUSSION

The use of CRISPR-Cas-based strategies to both cure or protect against invasion of AMR plasmids has been previously studied (Bikard et al., 2014; Chen et al., 2024; Citorik et al., 2014; Dong et al., 2019; Kim et al., 2016; Zhou et al., 2023). In this study, we showed that the mobile CRISPR-Cas9 plasmid pKJK5::csgc[*bla*_CTX-M-15_] protects an initially plasmid-free *E. coli* DH5α population when delivered either prophylactically before or simultaneously with exposure to *K. pneumoniae* carrying an AMR plasmid of clinical significance in an experimental community. Transfer levels below the limit of detection did not allow us to draw a conclusion of whether prophylactic or simultaneous exposure led to stronger protection against AMR plasmid dissemination; in both cases AMR plasmid dissemination into *E. coli* DH5α was reduced by at least ∼two orders of magnitude.

Unsurprisingly, the *E. coli* K12 host of the CRISPR-Cas9 plasmid showed strong protection against *K. pneumoniae* AMR plasmid pDSU25-1.1F uptake (***Supplementary figure 2***; and consistently observed across experiments). This strong protective effect of CRISPR-Cas9 against AMR plasmid uptake when it is already established in a host is in line with previous work (Bikard et al., 2014; Sünderhauf et al., 2023; Upreti et al., 2024), and reminiscent of a high, universal, benefit of CRISPR-Cas9 targeting while resident in its host cell. In contrast, mobile CRISPR-Cas9 directed against TA-carrying plasmids which are resident in a target host is less effective (Sünderhauf et al., 2026) and leads to cell death triggered by Cas9-induced plasmid loss (Citorik et al., 2014; Derollez et al., 2026).

Our experimental data was consistent with this phenomenon: *K. pneumoniae* readily took up non-targeting pKJK5::csgc[NT], but showed a marked decrease in uptake of pKJK5::csgc[*bla*_CTX-M-15_] (where CRISPR-Cas9 targets pDSU25-1.1F) (***Supplementary figure 3D*** and ***4D***). This low proportion of K. pneumoniae + pKJK5::csgc[*bla*_CTX-M-15_] transconjugants may be driven by a reduction of transconjugant viability during CRISPR-Cas9 targeting. Indeed, pDSU25-1.1F carries several TA systems with established post-segregational killing activity (***Table 1***; e.g. two tacAT systems (Fraikin & Van Melderen, 2024). More generally, TA systems are commonly found on *K. pneumoniae* plasmids (Wei et al., 2016) and other AMR plasmids including *bla*_CTX-M_-carrying IncF plasmids (Mnif et al., 2010).

This means that while protection of a CRISPR-Cas plasmid-carrying strain from AMR plasmid uptake is generally feasible and straightforward, curing of an already established AMR plasmid in a target host is not trivial. For this reason, we instead here highlight the ability of pKJK5::csgc to protect a naïve, plasmid-free focal recipient from AMR plasmid uptake while exposed to AMR and CRISPR-Cas9 plasmids simultaneously.

The presence of pKJK5::csgc in the synthetic community not only reduced AMR plasmid dissemination but also impacted community composition by reducing the density of the *K. pneumoniae* strain. Such *K. pneumoniae* reduction was independent from CRISPR-Cas9 immunity, since it was already observed in the pKJK5::csgc[NT] treatment. Therefore, reduction of the *K pneumoniae* burden might be attributed to a fitness cost of acquiring pKJK5::csgc. Consistent with this, high transfer of pKJK5::csgc into *K. pneumoniae* was observed in both experiments (***Supplementary figure 3D*** and ***4D***). The reduction of the *K. pneumoniae* burden was further maximised in presence of pKJK5::csgc[*bla*_CTX-M-15_], likely due to the combined effect between plasmid carriage fitness cost and TA-dependent post segregational killing after CRISPR-Cas9 cleavage.

Throughout our experiments, under liquid-broth mating conditions we were unable to observe any effects of pDSU25-1.1F transfer or *K. pneumoniae* reduction. This was due to low pKJK5::csgc transfer to both *E. coli* DH5α (∼10^-2^ transconjugants per recipients, ***Figure 1B***) and *K. pneumoniae* (<10^-3^ transconjugants per recipients, ***Supplementary figure 3D*** and ***4D***, liquid-broth). The inconsistency between mating conditions in the protective *E. coli* K12 + pKJK5::csgc[*bla*_CTX-M-15_] effect observed in the synthetic community highlights the importance of tailoring the CRISPR-Cas9 delivery vehicle to its specific application: IncP-based CRISPR-Cas9 tools such as pKJK5::csgc may be a good option when microbes on a surface are targeted, for instance within patients. Solid-surface mating has been described as a good approximation for mucosal and epithelial gut environments (Ott & Mellata, 2022). In contrast, when application conditions are closer to liquid-broth mating, the use of mobile plasmids from other families such as IncF can facilitate high transfer (Pulavan et al., 2026). These conditions are relevant to microbes in suspension, such as when preventing AMR plasmid transfer during a sewage spill, or indeed in other gut micro-niches (Ott & Mellata, 2022). In order to fully tailor a mobile CRISPR-Cas9 tool to its desired application, conjugation efficiency of the delivery vehicle should be tested in its relevant environment, while also considering other environmental factors such as pH and oxygen availability.

Overall, we showed that mobile CRISPR-Cas9 plasmid pKJK5::csgc[*bla*_CTX-M-15_] is a promising tool to prevent *K. pneumoniae* AMR plasmid transfer within a synthetic community. Importantly, protection is not limited to prophylactic delivery of CRISPR-Cas9 but can also be achieved when introducing the CRISPR-Cas9 host simultaneous to AMR plasmid host exposure. However, protection is only achieved under conditions that allow higher transfer of the CRISPR-Cas9 plasmid than of the AMR plasmid. These findings should be considered when applying CRISPR-Cas-based tools to prevent AMR dissemination.

## 4. METHODS

### 4.1 Genomic characterisation of clinical *K. pneumoniae* isolate

Clinical isolate *K. pneumoniae* MA48971.2 (BioProject PRJNA1550647) was obtained from a patient at University Medical Center, Utrecht (Netherlands) in 2018. We obtained long-and short-read sequencing from MicrobesNG sequencing service and assembled trimmed reads using Unicycler v0.5.1+galaxy0 (Wick et al., 2017) on the UseGalaxy EU server (Ostrovsky et al., 2022). The genome was annotated using Bakta v1.12.0 (Schwengers et al., 2021) and Kleborate v3.2.4 (Lam et al., 2021) was used to identify the sequence type (ST) of the isolate and screen for virulence and resistance genes. Plasmids were screened for replicon types and antimicrobial resistance genes using Abricate v1.4.0 (Seemann, 2026) with the PlasmidFinder (2026-Jul-7) (Carattoli et al., 2014) and ResFinder (2026-Apr-3) (Florensa et al., 2022) databases, respectively. Plasmid mobility was predicted using Mobsuite MOB-typer v3.1.9 (Robertson & Nash, 2018), and where the mobility prediction was non-mobile, sequences were additionally compared with putative oriT sequences (Ares-Arroyo et al., 2024) revealing putative origins of transfer to varying degrees of confidence across all plasmids. Defence systems were identified using DefenseFinder v2.0.0 (Tesson et al., 2022).

### 4.2 Strains, isolates, and plasmids

All strains, plasmids of interest, and selection regimes used are summarised in ***Table 2***. *E. coli* K12::mCherry (Klümper et al., 2015) has a chromosomal *aphA* Kanamycin resistance tag, alongside chromosomal *mCherry* and *lacI* insertion. This means K12::mCherry represses GFP expression of pKJK5::csgc, and appears red fluorescent only, allowing confident identification of transconjugants using fluorescence alongside selective plating. *E. coli* DH5α::SmR (Sünderhauf et al., 2026) is chromosomally tagged with an *aad* streptomycin resistance gene and derived from *E. coli* laboratory strain DH5α.

**Table 2:**
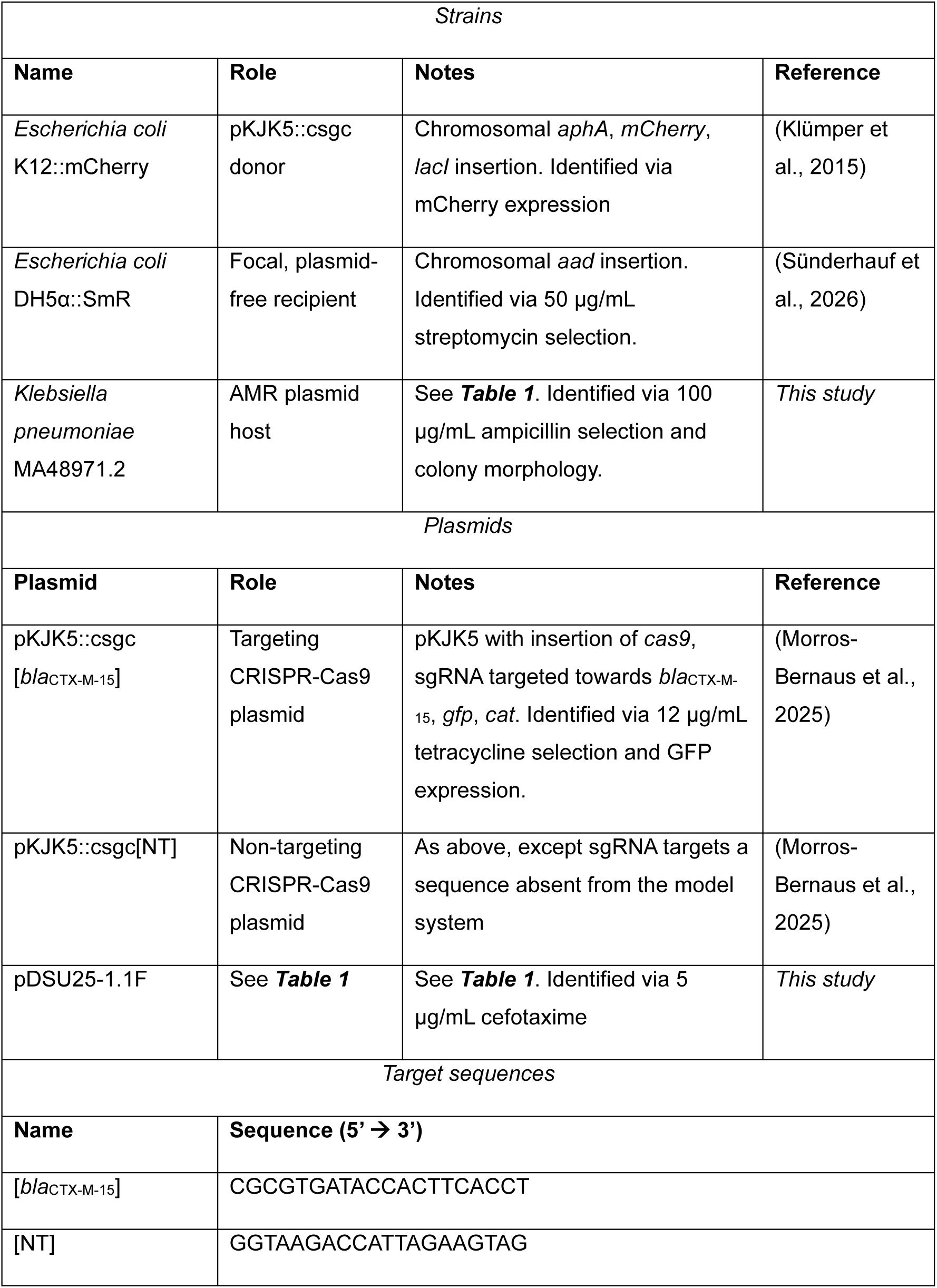
Strains and plasmids used in mating experiment.

pKJK5::csgc[*bla*_CTX-M-15_] and [NT] (Morros-Bernaus et al., 2025) are pKJK5 derivatives carrying Cas9, sgRNA, *gfp*, and *cat* chloramphenicol resistance inserted into pKJK5’s inactive *IntI1* gene. All inserted genes are under control of constitutive promoters; *gfp*’s promoter is in addition *lacI*-repressible and therefore inactive in *E. coli* K12::mCherry.

### 4.3 Mating experiments

Cultures were prepared by inoculating single colonies from a streak plate into 5 mL lysogeny broth (LB) supplemented with appropriate antibiotics to ensure plasmid maintenance, when needed (see ***Table 2***). These cultures were incubated overnight at 37°C at 180 rpm. *E. coli* cells (K12 and DH5α) were centrifuged at 3500 rpm for 10 minutes. The supernatants were discarded and pellets were washed twice in M9 buffer (22.4 mM Na2HPO4 7H2O, 22.0 mM KH2PO4, 8.6 mM NaCl, 18.7 mM Nh4Cl). *K. pneumoniae* cells were centrifuged as 5 x 1 mL aliquots at 14,800 rpm for 5 minutes, the supernatant was discarded, and the cells were washed in M9 buffer and transferred to a single receptacle. All strains were OD-adjusted to OD_600_ ∼ 0.5 with M9 buffer.

For one-to-one plasmid transfer experiments, two strains were used in each mating. In liquid-broth mating, 50 µL of each OD-adjusted strain was added to 5 mL LB and samples were incubated for 48 hours at 37°C at 50 rpm. For controls, M9 buffer was used in place of a second mating partner. The mixtures were vortexed for 5 seconds to stop conjugation. In solid-agar mating, 50 µL of each OD-adjusted strain (or M9 for controls) were centrifuged together at 14,800 rpm for 2 minutes, 70 µL of the supernatant was discarded, leaving a 30 µL volume for resuspension. Six 5 µL drops of the suspension of each biological replicate were plated on LB agar, allowed to dry, and incubated at 37°C for 48 hours. Cells were recovered by pipetting 1 mL of M9 buffer over the drops and resuspending with a hockey stick and pipette until the drops were no longer visible.

In the three-strain mating experiments where *E. coli* K12 + pKJK5::csgc was added prophylactically, the protocol described above was used with the following amendments. 50 µL of each OD-adjusted *E. coli* mating partner were incubated in either condition and recovered after two hours. In liquid-broth conditions, 50 µL of *K. pneumoniae* was added to the LB broth and incubation continued for 48 hours. In solid-agar conditions, 50 µL of OD-adjusted *K. pneumoniae* was added to ∼ 600 µL of recovered cells and centrifuged for 2 minutes. Supernatant was discarded leaving ∼ 30 µL to resuspend the cell pellet. Six 5 µL drops of suspension were plated onto LB agar, left to dry, and incubated for 48 hours. Cells in the solid-agar condition were recovered using 2 mL M9 buffer following the 48-hour incubation.

Where all three strains were mated simultaneously, 50 µL of each OD-adjusted strain was used to inoculate either liquid-broth or solid-agar medium as initially described. Incubation continued in both conditions uninterrupted over 48 hours.

Samples were taken and plated with selective medium after 48-(all) and 2-hour (only prophylactic mating) incubation periods. Antibiotics were used at selective concentrations listed in ***Table 2***, and colony identities were confirmed by colony morphology and fluorescence where relevant. *E. coli* DH5α + pDSU25-1.1F transconjugants were recovered applying selection for only pDSU25-1.1F, independent from pKJK5::csgc presence.

All experiments were performed with five biological replicates unless for the one-to-one mating between *K. pneumoniae* and *E. coli* DH5α, where three biological replicates were performed.

### 4.4 Data and statistical analysis

Data processing, visualisation and statistical analyses were performed in R v4.4.2 (R Core Team (2021)) using R studio 2023.12.1.402 (Posit team (2024)). Data processing and plotting were performed using the packages in *tidyverse* v2.0.0 (Wickham et al., 2019). For the three-strain community generated sequentially in the liquid-broth mating conditions, one replicate from the pKJK5::csgc[NT] and one from the pKJK5::csgc[*bla*_CTX-M-15_] were discarded when analysing the overall community composition since *E. coli* K12 density could not be measured due to the dilution used for plating, leading to N = 4. Additionally, for the three strains added simultaneously in liquid-broth mating conditions, one replicate from the no plasmid *E. coli* K12 treatment was discarded since no *E. coli* DH5α was inoculated. Additionally, another replicate from the same treatment was discarded to study the overall community composition since *E. coli* K12 density could not be measured due to the dilution used for plating, leading to N = 3.

Statistical modelling was performed using package *lme4* v1.1.37 (Bates et al., 2015) and *compositions* v2.0.8 (van den Boogaart & Tolosana-Delgado, 2008). Statistical significance of fixed effects was determined through comparing nested models using chi-squared tests (α = 0.05), starting from a global model containing all biologically relevant fixed effects and interaction terms. Model assumptions were evaluated using the R package *DHARMa* v0.4.7 (Hartig, 2026). Post-hoc analyses were conducted using the *emmeans* package v1.11.1 (Searle et al., 1980). Pairwise comparisons were adjusted using Tukeýs correction.

Across experiments, the conjugation efficiencies of the different plasmids were modelled using gamma family generalised linear models (GLMs) with a log link function. Due to large differences in transconjugant abundance among the different plasmid treatments, concentrations of recipients and of transconjugants (CFU/mL) were logarithmically transformed. Therefore, for all models log10 (transconjugants) / log10 (recipients) was used as a response variable. When no transconjugants were recovered the limit of detection, set to a theoretical recovery of 0.5 transconjugants, was used instead. Unless otherwise specified, all fitted models met the underlying assumptions, with diagnostic checks indicating no evidence of residual deviations, over-or under dispersion, influential outliers or heteroscedasticity.

For the 1:1 mating datasets, separate models were fitted for pDSU25-1.1F (*Figure 1A*) and pKJK5::csgc[NT] (*Figure 1B*) transfer, respectively. In both cases, mating condition was included as explanatory variable which had a significant effect on the response variable (Gamma GLM, χ² = 0.27, *p* <0.0001 and Gamma GLM, χ² = 0.07, *p* <0.0001). For the sequential three-strain synthetic community dataset, four models were fitted to assess conjugation efficiencies of (i) pDSU25-1.1F into *E. coli* DH5α (*Figure 2B* and *Supp.Fig. 3A*), (ii) pKJK5 into *E. coli* DH5α during the first 2 hours of mating (*Supp. Fig. 1*), (iii) pKJK5 into *K. pneumoniae* (*Supp. Fig. 3D*) and (iv) pDSU25-1.1F transfer into *E. coli* K12 + pKJK5::csgc (*Supp. Fig. 2*). Models (i), (ii) and (iii) included mating condition and plasmid treatment as explanatory variables, as well as their interaction term due to its biological relevance. Besides, in all cases, the interaction between mating condition and plasmid treatment had a significant effect on the response variable ((i) Gamma GLM, χ² = 0.54, *p* <0.0001; (ii) Gamma GLM, χ² = 0.16, *p* <0.0001 and (iii) Gamma GLM, χ² = 0.15, *p* <0.0001). Model (iv) only included plasmid treatment as explanatory variable which had a significant effect on the response variable (Gamma GLM, χ² = 1.10, *p* <0.0001). For pDSU25-1.1F transfer into *E. coli* DH5α (i), model validation revealed an influential outlier corresponding to one replicate from the no plasmid treatment under solid-agar mating conditions, where no transconjugants were recovered (triangle replicate in the no plasmid treatment in *Figure 2B*). Model optimization was performed after removing this outlier. For pKJK5 transfer into *E. coli* DH5 (ii), model validation revealed under-dispersion and influential outliers corresponding to the liquid-broth mating replicates from [NT] and [*bla*_CTX-M-15_] where no transconjugants were recovered (triangle replicates under liquid-broth mating in *Supp. Fig. 1*). Model optimization was performed after removal of these outliers. For pDSU25-1.1F transfer into *E. coli* K12 + pKJK5::csgc (iv) model validation revealed an influential outlier corresponding to one replicate in the pKJK5::csgc[*bla*_CTX-M-15_] treatment with higher limit of detection. Sensitivity analysis excluding this observation yielded the same conclusion regarding the significance of the Plasmid (*p*<0.0001 compared with *p*<0.0001 in the full dataset). Therefore, the observation was maintained in the dataset since there was no biological reason for exclusion. For the simultaneous three-strain synthetic community dataset, two models were also fitted to assess conjugation efficiencies of (i) pDSU25-1.1F into *E. coli* DH5α (*Figure 3A* and *Supp.Fig. 4A*) and (ii) pKJK5 into *K. pneumoniae* (*Supp. Fig. 4D*). The two models included mating condition and plasmid treatment as explanatory variables, as well as their interaction term due to its biological relevance. Besides, in all cases, the interaction between mating condition and plasmid treatment had a significant effect on the response variable (Gamma GLM, χ² = 0.71, *p* <0.0001 and Gamma GLM, χ² = 0.77, *p* <0.0001, respectively). For pDSU25-1.1 transfer into *E. coli* DH5α (i), model validation revealed an influential outlier attributed to a replicate in the No plasmid treatment with lower transconjugants in liquid-broth mating. Sensitivity analysis excluding this observation yielded the same conclusion regarding the significance of the Bacteria × Plasmid interaction (*p*<0.0001 compared with *p*<0.0001 in the full dataset). Therefore, the observation was maintained in the dataset since there was no biological reason for exclusion.

The bacterial proportions of the sequential (*Figure 2C* and *Supplementary figure 3B*) and simultaneous (*Figure 3B* and *Supplementary figure 4B*) synthetic communities were assessed using compositional data analysis. Relative abundances of *K. pneumoniae, E. coli* DH5α and *E. coli* K12 were transformed using the isometric log-ratio (ILR) transformation, and the effect of plasmid treatment on bacterial community was tested using a multivariate analysis of variance (MANOVA) with Pillaís trace (V).

The overall cell densities (CFU/mL) of the sequential (*Figure 2D* and *Supplementary figure 3C*) and simultaneous (*Figure 3C* and *Supplementary figure 4C*) synthetic communities were modelled using gamma family generalised linear models (GLMs) with a log link function. Cell densities were logarithmically transformed. For each community, models for solid-agar and liquid-broth mating datasets were fitted individually. Bacteria type, plasmid treatment and their interaction term had a significant effect on the response variable (Sequential bacterial community solid-agar mating: Gamma GLM, χ² = 0.009, *p* = 0.008; Simultaneous bacterial community solid-agar mating: Gamma GLM, χ² = 0.007, *p* = 0.001 and Simultaneous bacterial community liquid-broth mating: Gamma GLM, χ² = 0.03, *p* = 0.001). For the sequential bacterial community and simultaneous bacterial community solid-agar mating models, model diagnostics identified one potentially influential observation in each experiment. Because there was no experimental or biological justification for excluding these observations, they were retained in the primary analyses. Sensitivity analyses excluding these observations yielded the same conclusion regarding the significance of the Bacteria × Plasmid interaction (sequential experiment: *p* = 0.0026 compared with *p* = 0.008 in the full dataset; simultaneous experiment: *p* = 0.006 compared with *p* = 0.001 in the full dataset). For the sequential bacterial community liquid-broth mating the interaction term had no significant effect (Gamma GLM, χ² = 0.002, *p* = 0.76). However, the interaction remained in the model due to its biological relevance.

### 4.5 DATA AVAILABILITY

The raw data together with the R project containing all the scripts used to generate the plots and to analyse the data statistically can be found in the supplementary excel files and R project deposited in GitHub: https://github.com/claudiamb98/CRISPR-Cas9-treatment-Klebsiella-pneumoniae.

The assembled genome together with the raw sequencing data of the clinical *K. pneumoniae* MA48971.2 is deposited in the NCBI BioProject: PRJNA1550647 under the BioSample: SAMN64440691.

## Supporting information

Raw data and data analysis

## ACKNOWLEDGMENTS

We thank Dr Fernanda Paganelli (UMC Utrecht, The Netherlands) for supplying clinical K. pneumoniae isolate MA48971.2 and Dr Elizabeth Pursey for providing preliminary sequencing data and analyses.

## SUPPLEMENTARY FIGURES

**Supplementary figure 1:**
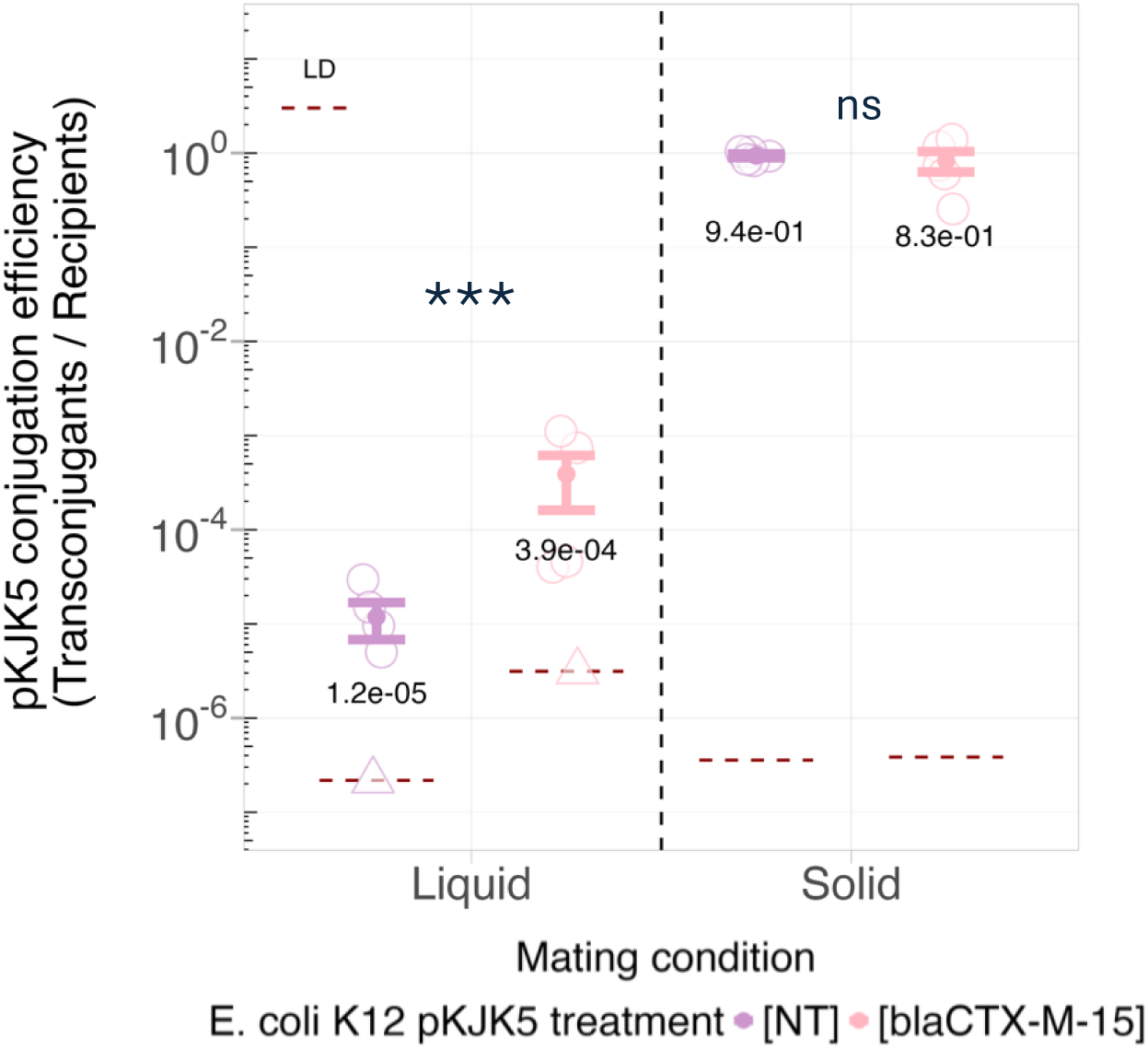
Transfer of pKJK5::csgc into *E. coli* DH5α after the initial two-hour mating between *E. coli* K12 + pKJK5::csgc[NT] or pKJK5::csgc[*bla*_CTX-M-15_], before adding *K. pneumoniae* into the synthetic community. Liquid-broth and solid-agar mating were performed. Plasmid transfer is shown as conjugation efficiency, which is calculated as transconjugants (*E. coli* DH5α + pKJK5::csgc) divided by recipients (overall *E. coli* DH5α population). Mean and standard error are shown as point and bars and individual replicates are plotted as circles. In those cases where pKJK5-carrying transconjugants were not observed the limit of detection (LD) of the assay is plotted instead (denoted by triangles). LD was determined based on the plating volume and dilution factors and was set at 0.5 transconjugants detected in the non-diluted sample when transconjugants were detected, or in the plated dilution when no transconjugants were detected. Since LD is different for each individual replicate, the highest LD within treatment is plotted as a dashed red line. *** *p* = 0.0002; gamma GLM, emmeans contrast Tukey-adjusted; n.s. – not significant.

**Supplementary figure 2:**
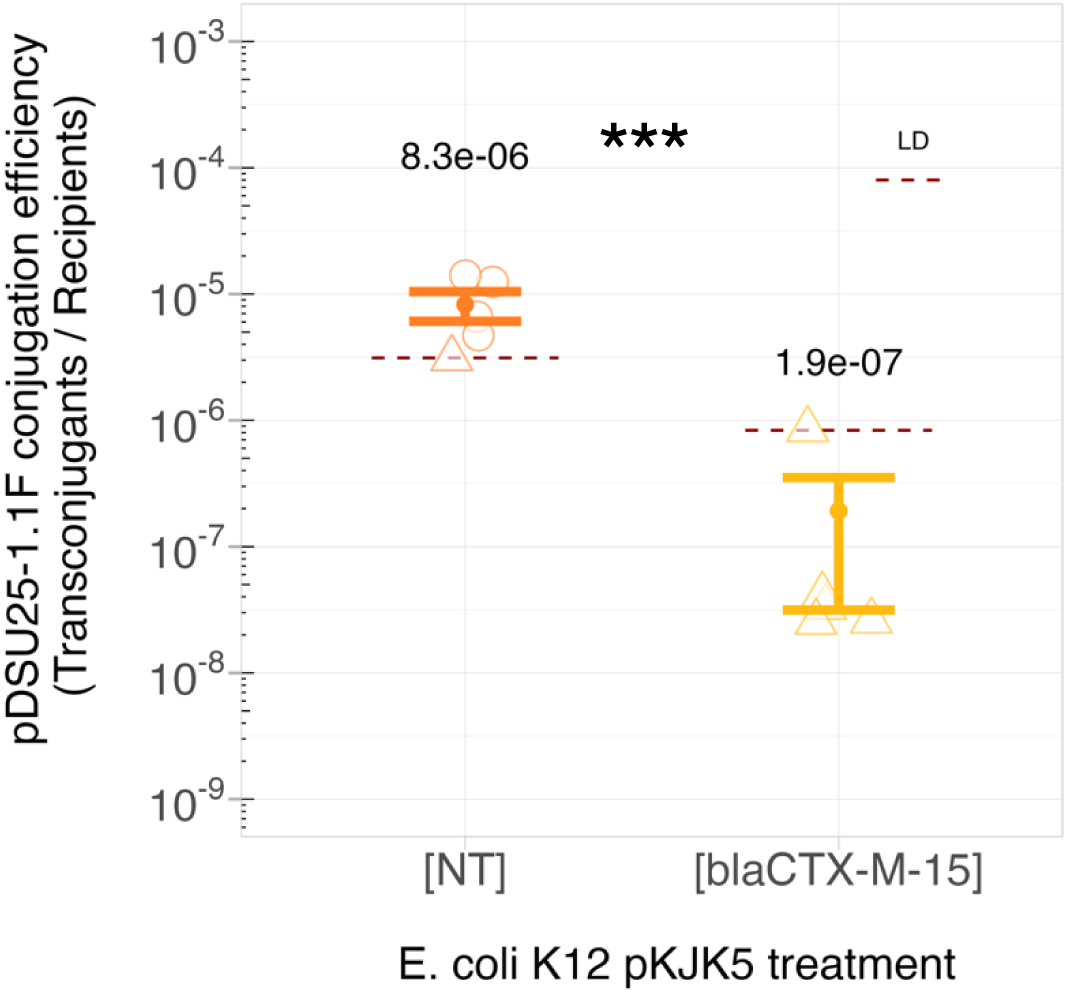
CRISPR-Cas9 immunity (pKJK5::csgc[*bla*_CTX-M-15_]) against uptake of a *bla*_CTX-M-15_-carrying *K. pneumoniae* plasmid (pDSU25-1.1F) in the *E. coli* K12 host, in the three-strain sequential synthetic community (Figure 2A) undergoing solid-agar mating. Plasmid transfer is shown as conjugation efficiency, which is calculated as transconjugants (*E. coli* K12 + pKJK5::csgc + pDSU25-1.1F) divided by recipients (overall *E. coli* K12 + pKJK5::csgc population). Mean and standard error are shown as point and bars and individual replicates are plotted as circles. In those cases where pDSU25-1.1F-carrying transconjugants were not observed, the limit of detection (LD) of the assay is plotted instead (denoted by triangles). LD was determined based on plating volume and dilution factors and was set to 0.5 transconjugants detected in the non-diluted sample. Since LD is different for each individual replicate, the highest LD within treatment is plotted as a dashed red line. *** *p* = 0.0002; gamma GLM, emmeans contrast Tukey-adjusted.

**Supplementary figure 3:**
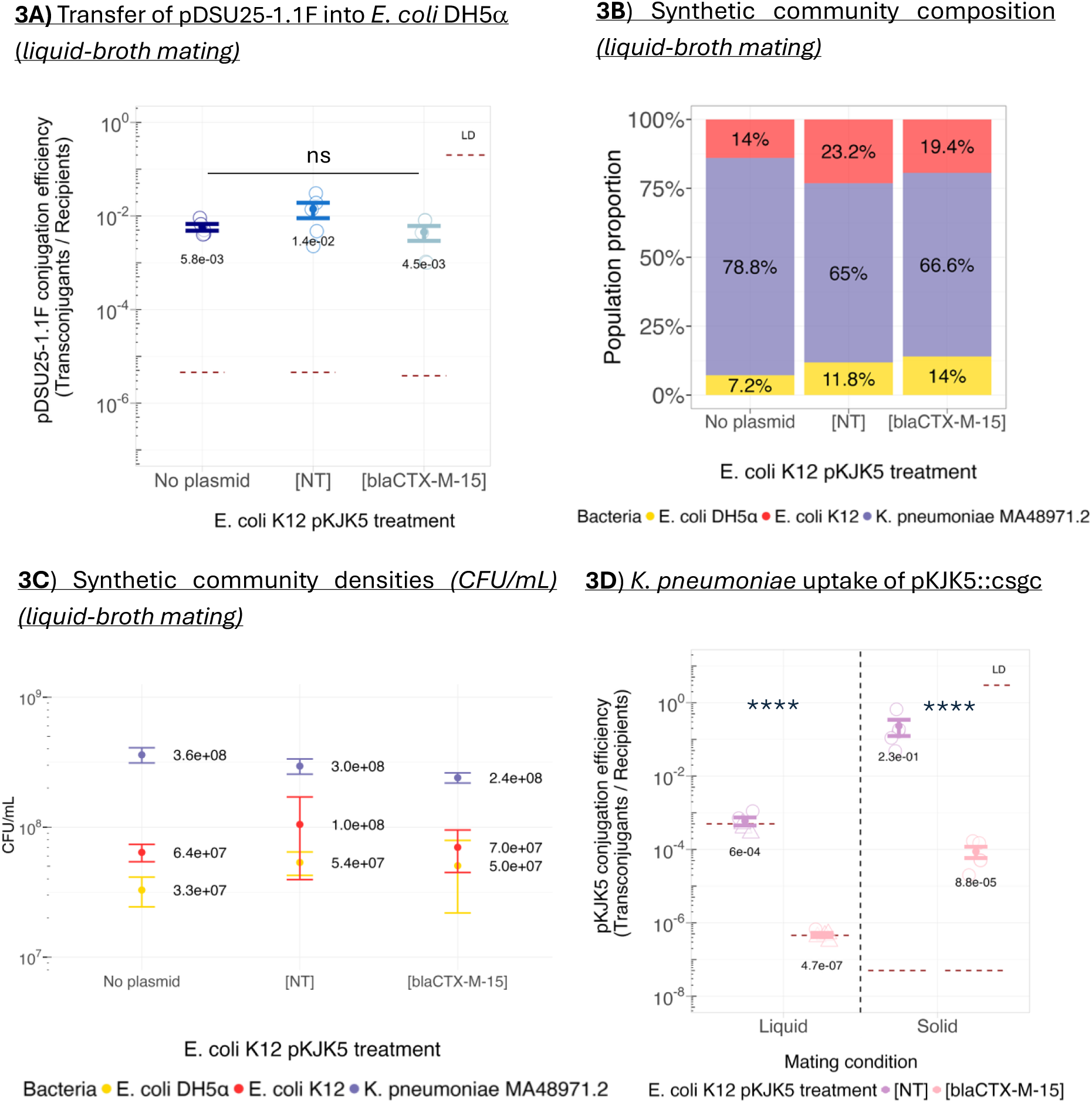
Supplementary figure 3 shows data from the sequential three-strain synthetic community. Liquid-broth mating results from the sequential three-strain synthetic community (*E. coli* K12, *E. coli* DH5α and *Klebsiella pneumoniae* strain MA48971.2; Figure 2). **A)** Transfer of pDSU25-1.1F into *E. coli* DH5α shown as conjugation efficiency, which is calculated as transconjugants (*E. coli* DH5α + pDSU25-1.1F) divided by recipients (overall *E. coli* DH5α population). Mean and standard error are shown as point and bars, individual replicates are plotted as circles. N.s. not-significant as assessed by gamma GLM, emmeans contrast Tukey-adjusted. **B-C)** Synthetic community composition (**B**) and bacterial densities (**C**) across *E. coli* K12 treatments. *E. coli* K12 is shown in red, *E. coli* DH5α in yellow and *K. pneumoniae* in purple. Means from 5 replicates are plotted across treatments apart from one replicate from the [NT] treatment and one from the [*bla*_CTX-M-15_] treatment being excluded because *E. coli* K-12 density was not measured, due to the dilution used for plating. Discarding datapoints did not affect the overall population proportions or densities. **B**) Synthetic community composition in liquid-broth mating. Plasmid treatment did not significantly affect bacterial community composition (MANOVA, V = 0.23, F (4,20) = 0.65, *p* = 0.6). **C)** Bacterial density of the respective strains (CFU/mL) in liquid-broth mating. Mean and standard error are shown as point and bars. No significant differences within strains were detected (gamma GLM, emmeans contrast Tukey-adjusted, all *p* > 0.05)**. D)** *K. pneumoniae* uptake of pKJK5::csgc in both liquid-broth and solid-agar mating. Plasmid transfer is shown as conjugation efficiency, which is calculated as transconjugants (*K. pneumoniae +* pKJK5::csgc) divided by recipients (overall *K. pneumoniae* population). \*\*\*\**p*<0.0001; gamma GLM, emmeans contrast Tukey-adjusted *p*<0.0001. Mean and standard error are shown as point and bars and individual replicates are plotted as circles. For those cases where pKJK5-carrying transconjugants were not observed, the limit of detection (LD) of the assay is plotted instead (denoted by triangles). LD was determined based on the plating volume and dilution factors and was set at 0.5 transconjugants detected in the non-diluted sample when transconjugants were detected, or in the plated dilution when no transconjugants were detected. Since LD is different for each individual replicate, the highest LD within treatment is plotted as a dashed red line.

**Supplementary figure 4:**
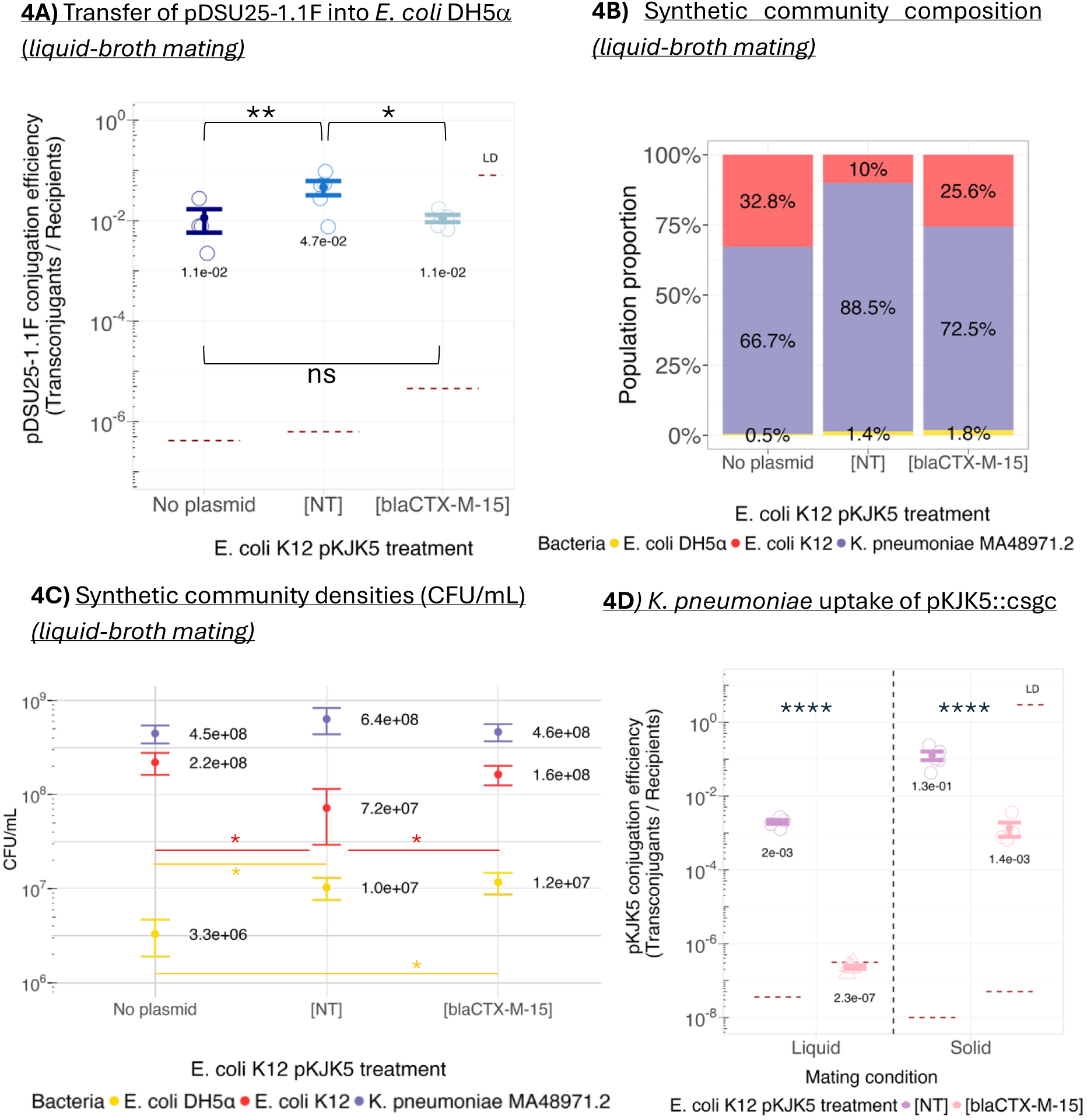
Supplementary figure 4 shows data from the simultaneous three-strain synthetic community. Liquid-broth mating results from the three-strain synthetic community (*E. coli* K12, *E. coli* DH5α and *Klebsiella pneumoniae* strain MA48971.2, as in Figure 2 except all strains were added simultaneously). **A)** Transfer of the *K. pneumoniae* plasmid pDSU25-1.1F into *E. coli* DH5α. Plasmid transfer is shown as conjugation efficiency, which is calculated as transconjugants (*E. coli* DH5α + pDSU25-1.1F) divided by recipients (overall *E. coli* DH5α population). Mean and standard error are shown as point and bars and individual replicates are plotted as circles. Means from 5 replicates are plotted across treatments unless for the “no plasmid” where one replicate was discarded since no *E. coli* DH5α was detected. N.s. not significant; * *p* = 0.03, ** *p* = 0.005 as assessed by gamma GLM, emmeans contrast Tukey-adjusted. **B-C)** Synthetic community composition (**B**) and bacterial densities (**C**) across *E. coli* K12 treatments. *E. coli* K12 is shown in red, *E. coli* DH5α in yellow and *K. pneumoniae* in purple. **B)** Synthetic community composition in liquid-broth mating. Means from 5 replicates are plotted across treatments unless for “no plasmid” where two replicates were discarded due to either no *E. coli* DH5α or lack of *E. coli* K12 density measures due to the dilution used for plating. Plasmid treatment significantly affected bacterial community composition (MANOVA, V = 0.78, *F* (4,20) = 3.19, *p* = 0.034). **C)** Bacterial density of the respective strains (CFU/mL), mean and standard error are shown as point and bars. \**p*<0.05 as assessed by gamma GLM, emmeans contrast Tukey-adjusted. All other within-strain comparisons remained non-significant. **D)** Transfer of the pKJK5::csgc[NT] or pKJK5::csgc[*bla*_CTX-M-15_] from *E. coli* K12 to the *K. pneumoniae* strain in both liquid-broth and solid-agar mating. Plasmid transfer is shown as conjugation efficiency, which is calculated as transconjugants (*K. pneumoniae +* pKJK5::csgc) divided by recipients (overall *K. pneumoniae* population). Mean and standard error are shown as point and bars and individual replicates are plotted as circles. For those cases where pKJK5-carrying transconjugants were not observed, the limit of detection (LD) of the assay is plotted instead (denoted by triangles). LD was determined based on the plating volume and dilution factors and was set at 0.5 transconjugants detected in the non-diluted sample when transconjugants were detected, or in the plated dilution when no transconjugants were detected. Since LD is different for each individual replicate, the highest LD within treatment is plotted as a dashed red line. **** *p*<0.0001; gamma GLM, emmeans contrast Tukey-adjusted *p*<0.0001.

