## Supplementary material for "Prevention of *Klebsiella pneumoniae* antibiotic resistance plasmid dissemination by prophylactic or simultaneous CRISPR-Cas9 treatment": Raw data and data analysis: Readme.docx

Manuscript title:

Description:

The DATA folder contains an R project with excel files containing the raw data from the different experiments, along with the corresponding R scripts used for data analysis. Additionally, a README file is included in the R project, providing instructions on how to access and run the datasets and scripts.
